# Transcriptomic Profiling of Thyroid Eye Disease Orbital Fibroblasts Identifies Sorafenib as a Novel Therapeutic

**DOI:** 10.64898/2026.04.21.719973

**Authors:** Kyle Yuan, Phillip Truong, Charkira C. Patrick, Emma Ushchak, Elisa Roztocil, Steven E. Feldon, Collynn F. Woeller

## Abstract

Thyroid eye disease (TED) is a debilitating condition characterized by orbital fibroblast (OF) activation and excessive hyaluronic acid (HA) accumulation within the retro-ocular space. While IGF-1R blockade with teprotumumab has significantly advanced TED management, incomplete clinical responses and disease relapse underscore the need to identify alternative targets. In this study, we used high-throughput RNA sequencing to map the transcriptomic landscape in TED OFs compared with non-TED OF controls. Our analysis identified robust enrichment of pathways critical to the TED phenotype, including PI3K-AKT signaling, the platelet-derived growth factor (PDGF) pathway, and extracellular matrix remodeling. We validated several key upregulated mediators that may contribute to orbital remodeling, including FOXC2, HGF, MET, and HMGA2, alongside the downregulation of the Wnt antagonist SFRP2. By employing a computational drug-repositioning approach, we identified the multi-kinase inhibitor sorafenib, which targets VEGFR, PDGFR, and RAF, as a potent candidate to neutralize the TED-specific gene signature. Functional assays demonstrated that sorafenib dose-dependently inhibited PDGF-induced AKT phosphorylation and significantly attenuated HA synthesis in primary TED OFs. These results define a persistent, receptor tyrosine kinase-driven program in the TED orbit and suggest that multi-kinase inhibition represents a viable therapeutic strategy for refractory TED.

**Highlights:**

- Thyroid eye disease (TED) orbital fibroblasts exhibit a transcriptomic signature characterized by elevated PI3K/AKT, angiogenic, and growth factor signaling.
- Computational drug prediction identifies sorafenib as a candidate to reverse the TED gene signature.
- Sorafenib dose-dependently inhibits AKT activation and hyaluronic acid production in TED orbital fibroblasts.

## Introduction

Thyroid eye disease (TED) is a debilitating autoimmune condition characterized by expansion of orbital adipose tissue and fibrosis of the extraocular muscles, leading to proptosis, diplopia, and vision loss. The annual incidence of TED is 2 in every 10,000 individuals and is highly associated with Graves’ disease (GD): 25%-40% of individuals with GD are diagnosed with TED (1, 2). Among those, moderate-to-severe TED affects an estimated 4.4-9.0 per 100,000 individuals, with women disproportionately affected (3). Treatments for TED include systemic glucocorticoids, mycophenolate, and orbital decompression surgery (4–6), with systemic glucocorticoids as the mainstay of treatment (7–9). Teprotumumab, an insulin-like growth factor-1 receptor (IGF-1R) blocking antibody, has shown great promise in the US, with approximately 80% of patients experiencing a reduction in symptomatic disease (10). Clinical trials have demonstrated significant improvements in key clinical outcomes, including reductions in proptosis (≥2 mm), the Clinical Activity Score (CAS), diplopia, and quality-of-life scores (10, 11).

Despite these benefits, teprotumumab can cause clinically significant adverse events, including hearing loss, muscle spasms, inflammatory bowel disease flares, and hyperglycemia. In a study of 131 patients who received ≥4 infusions, 8.4% experienced severe adverse events and 12.2% discontinued therapy (12). In addition, 20%-30% of TED patients are refractory to first-line treatments, and 10%-20% relapse after treatment is withdrawn (7). The limited therapeutic options and variable response profiles have prompted the use of genomic approaches, such as RNA sequencing (RNA-seq), to better define TED pathogenesis at the molecular level.

Orbital fibroblasts (OFs) are central to TED pathogenesis because they can differentiate into adipocytes or myofibroblasts, secrete large amounts of hyaluronic acid (HA), and proliferate excessively (2, 13–15). TED OFs express both the thyroid-stimulating hormone receptor (TSHR) and IGF-1R(15–19). Stimulatory TSHR autoantibodies found in TED/Graves’ patients can promote adipogenesis, inflammation, and HA synthesis(20, 21). TSHR and IGF-1R converge on the phosphatidylinositol 3-kinase (PI3K)/AKT pathway, which is increasingly recognized as a nodal regulator of proliferation, cytokine production, and extracellular matrix (ECM) remodeling in TED (15, 22, 23). *In vivo*, TED OFs are also exposed to inflammatory mediators and additional receptor tyrosine kinase (RTK) ligands, including PDGF, VEGF, and TGFβ, which can further engage the PI3K/AKT axis and contribute to orbital pathology (24–28). The complex, dynamic interplay of these signals may explain the clinical heterogeneity observed in TED, in which some patients exhibit predominantly fat expansion, others extraocular muscle enlargement, and many present with a heterogeneous phenotype characterized by increases in both fat and muscle.

To identify new therapeutic targets in TED, and given that current animal models only partially reproduce the orbital phenotype (29–32), we utilized primary OFs from orbital fat decompression surgery, a physiologically relevant and widely validated system for dissecting TED-specific signaling (14, 33). Here, we performed transcriptome-wide expression profiling of primary TED and control OFs and applied gene ontology tools, including Enrichr (34–36), ToppGene (37–39), and DAVID (40, 41) to define dysregulated pathways in TED. We then used the LINCS L1000CDS² platform (42) to screen for FDA-approved therapeutics predicted to reverse the TED OF expression signature. This integrative strategy identified sorafenib, a multi-kinase inhibitor targeting RAF, VEGFR, and PDGFR, which we validated functionally in PDGFβ-stimulated TED OFs. Sorafenib dose-dependently suppressed TED-specific gene expression and inhibited HA production, demonstrating its therapeutic potential in TED.

## Results

### Transcriptomic Profiling Reveals Distinct Gene Expression Signatures in TED Orbital Fibroblasts

To define disease-associated transcriptional changes in TED, we performed RNA-seq on OFs from TED and non-TED donors. Differential expression analysis identified a robust set of upregulated and downregulated genes in TED OFs compared to non-TED controls **(Fig. 1A)**. Several of the most significantly altered genes are known regulators of extracellular matrix (ECM) remodeling, growth factor signaling, and fibroblast activation. A heatmap showing unsupervised hierarchical clustering of the top differentially expressed genes revealed strong clustering by disease state, demonstrating that the observed gene expression changes were consistent across individual donors **(Fig. 1B)**.

**Fig. 1.**
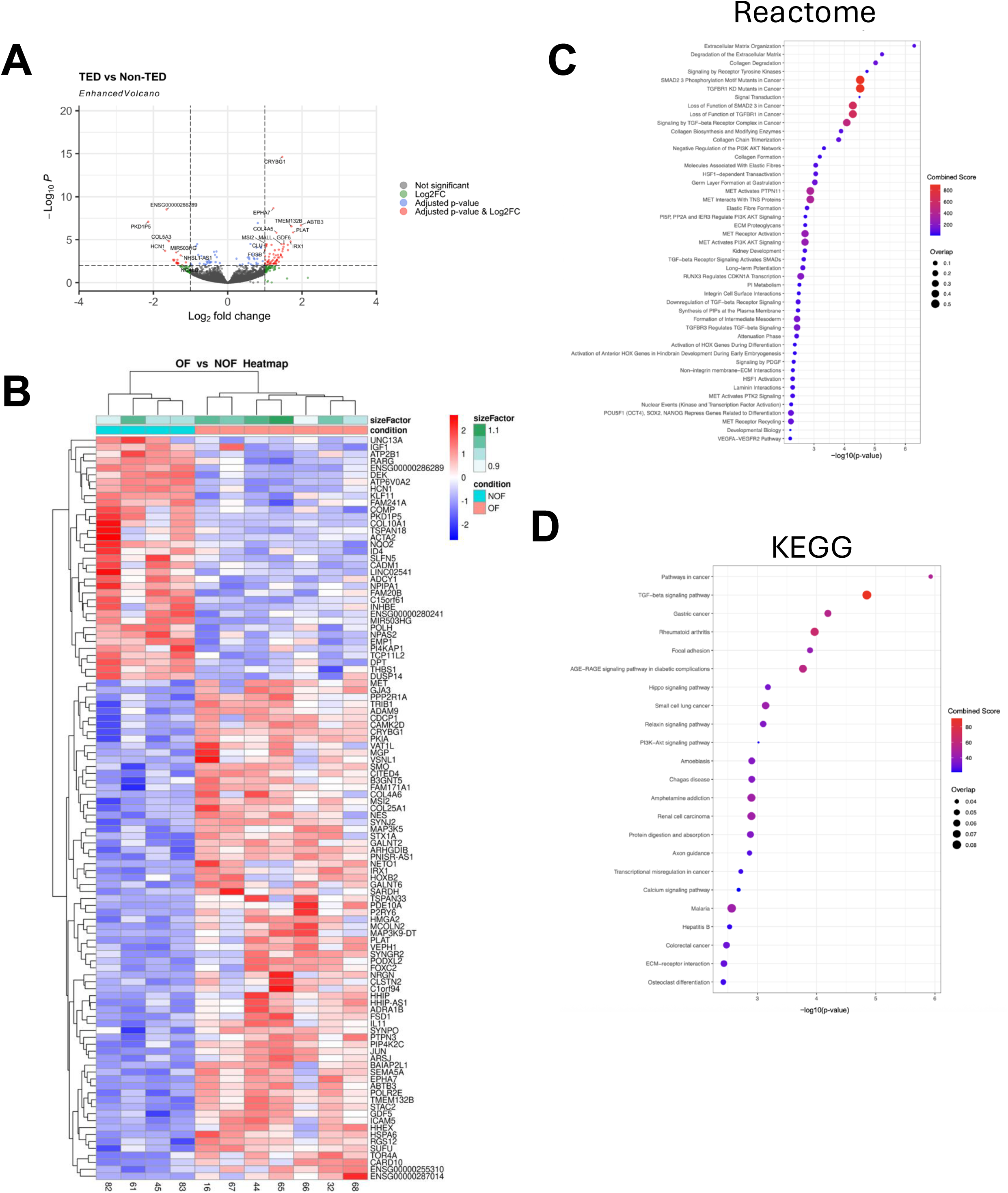
Differential gene expression analysis of TED versus non-TED orbital fibroblasts. (A) Volcano plot showing RNAseq results comparing TED (n = 7) and non-TED (n = 4) orbital fibroblasts. The x-axis represents log₂ fold change, and the y-axis represents –log₁₀ adjusted p-value. Genes meeting both the fold-change (log₂ FC > threshold) and adjusted p < 0.05 criteria are highlighted in red. (B) A heatmap of the top differentially expressed genes, ranked by adjusted p-value, demonstrates apparent clustering of TED and non-TED fibroblast strains. Expression values are shown as red (upregulated) or blue (downregulated) relative to the group mean pathway enrichment analysis of differentially expressed genes in TED OFs. Bubble plots display significantly enriched pathways identified by Enrichr using Reactome (C) (p-adj < 0.10) and KEGG (D) (p-adj < 0.05) databases. Upregulated genes were enriched in extracellular matrix (ECM) organization, collagen metabolism, receptor tyrosine kinase signaling, and TGFβ-related signaling. Bubble size indicates the proportion of input genes in each term; color represents the combined enrichment score.

### TED OFs are Enriched for PI3K/AKT and Angiogenic Signatures Compared to Non-TED OFs

A comparison of TED versus non-TED OFs identified 233 upregulated genes and 129 downregulated genes in TED (adjusted p < 0.05). Genes enriched in TED OFs included *HGF, MET, FOXC2, HMGA2, PLEKHG5, and JUN*. Gene Ontology (GO) analysis revealed activation of *PI3K /AKT signaling* (adjusted p = 0.018). Interestingly, *negative regulation of the PI3K/AKT network* (adjusted p = 0.027) was also significant. Two additional downstream pathways narrowly missed the significance cutoff, including *MET activating PI3K/AKT signaling* (adjusted p = 0.061) and *PI5P, PP2A, and IER3 regulating PI3K/AKT signaling* (adjusted p = 0.061) (**Fig. 1C, 1D, and Supplemental Table 1).**

TED OFs also showed significant enrichment of angiogenesis-associated pathways. These pathways included: *vascular smooth muscle cell/pericyte differentiation and proliferation* (adjusted p = 0.00008), *extracellular matrix organization* (adjusted p = 0.0004), *positive regulation of smooth muscle cell proliferation* (adjusted p = 0.023), *vascular endothelial cell activation by growth factors* (adjusted p = 0.037), and *sprouting angiogenesis* (adjusted p = 0.039). Upregulation of *SMAD3*, *TGFB1*, and *TGFBR2* contributed to these pathways, suggesting that the TGFβ signaling cascade plays a significant role in regulating angiogenesis **(Supplemental Table 1)**. BMP4 and SEMA5A were also upregulated in select angiogenic pathways. BMP4 is a well-established member of the TGFβ superfamily (43), and SEMA5A has previously been associated with angiogenic processes (44).

The extracellular matrix, as well as the synthesis and regulation of collagen, play essential roles in angiogenesis by forming a structural scaffold that supports endothelial cell migration (45). Pathways involving the extracellular matrix and collagen were significantly upregulated in TED. These pathways included *extracellular matrix organization* (adjusted p = 0.0004), *collagen degradation* (adjusted p = 0.002), and *collagen biosynthesis and modifying enzymes* (adjusted p = 0.009). Upregulated genes contributing to these pathways included collagen genes such as COL4A6, COL4A5, COL9A3, COL13A1, and COL25A1, as well as genes encoding metalloproteinases, including ADAMTS8, ADAM9, and MMP1 **(Fig. 1C, D, Supplementary Figure 1).** Significant pathways were identified among downregulated genes in TED OFs. However, these pathways lacked a consistent functional theme.

To validate our RNA-seq results, we performed RT-qPCR on a subset of differentially expressed genes. First, we assessed pro-angiogenic/RTK-linked genes identified in the transcriptomic analysis (*FOXC2, MET, HGF*). We confirmed that all were significantly higher in TED OFs than non-TED OFs (**Fig. 2A-C**). Second, we measured fibroblast activation/motility-associated transcriptional regulators (*HMGA2, PLEKHG5*), which were also elevated in TED OFs, consistent with an activated fibro-proliferative phenotype(**Fig. 2D and E**). Finally, we examined *SFRP2*, a negative regulator of Wnt signaling, which was reduced in TED OFs by RNAseq and confirmed to be significantly downregulated by qPCR (**Fig. 2F**). Together, these data validate the RNA-seq findings at the single-gene level and show that TED OFs upregulate angiogenic/RTK and fibroblast activation programs while downregulating the Wnt regulator, *SFRP2*.

**Fig. 2.**
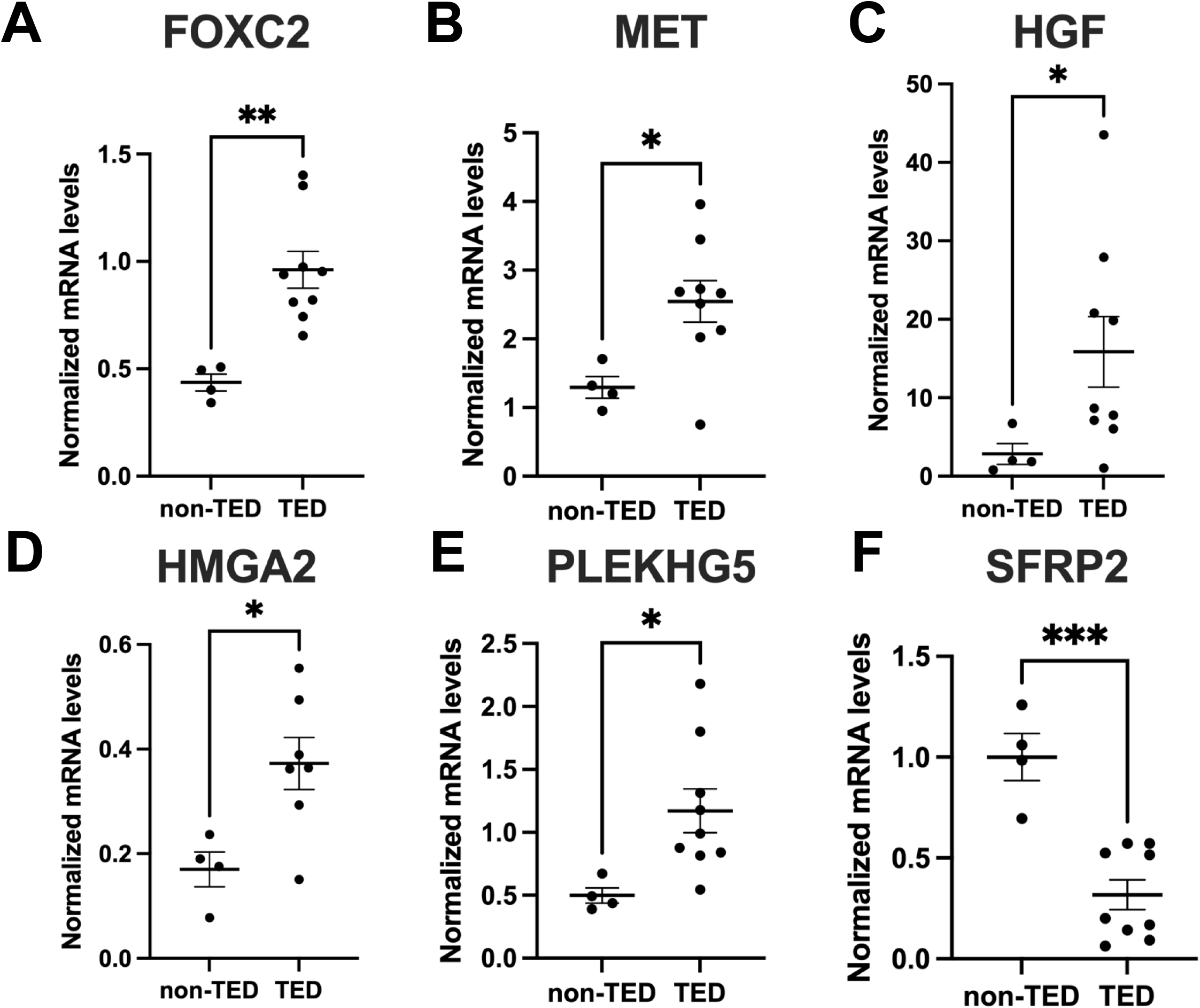
Validation of selected differentially expressed genes by RT-qPCR. Orbital fibroblasts from non-TED (n = 4) and TED (n = 7) donors were analyzed for: (i) angiogenic/RTK-associated genes, *FOXC2, MET, and HGF* (A, B, and C, respectively); (ii) fibroblast activation/motility regulators, *HMGA2, and PLEKHG5 (D and E)*; and (iii) the Wnt antagonist *SFRP2* (F). mRNA levels were normalized to *TBP* and *18S rRNA*. TED OFs showed significantly increased expression of *HGF, MET, FOXC2, HMGA2*, and *PLEKHG5*, and decreased expression of *SFRP2* compared with non-TED OFs. Data are mean ± SEM; p < 0.05, *p < 0.01, **p < 0.001 by unpaired two-tailed Student’s t-test.

### Connectivity Map Analysis Identifies Sorafenib as a Predicted Therapeutic Candidate

We next leveraged the LINCS L1000 Connectivity Map to identify compounds that could reverse the TED gene expression signature (46, 47). Compounds with strong negative connectivity scores are predicted to counteract the TED transcriptional program. Among FDA-approved drugs, mirdametinib, selumetinib, and trametinib (all MEK1/2 inhibitors), and sorafenib (a multikinase inhibitor targeting RAF, RET, VEGFR, and PDGFR) emerged as top hits **(Table 3)**.

We selected sorafenib for functional validation based on several considerations. First, sorafenib inhibits RAF (upstream of MEK1/2), providing broader pathway coverage than MEK-specific inhibitors. Second, sorafenib’s multi-target profile, including VEGFR and PDGFR, directly addresses the angiogenesis and PDGF signaling pathways enriched in the TED OF transcriptome. Third, sorafenib has demonstrated efficacy in other fibrotic disorders (48, 49). We therefore evaluated sorafenib’s capacity to suppress PDGFβ-induced activation in primary TED OFs.

### Sorafenib Suppresses PDGFβ-Induced Pathogenic Gene Expression in TED Orbital Fibroblasts

Since several of the enriched pathways involved receptor tyrosine kinases, including PDGF, VEGF, and MET receptors, we investigated whether the *in silico* predicted inhibitor, sorafenib, could functionally reverse this transcriptional state. First, we evaluated sorafenib’s ability to block PDGFβ-driven gene expression in TED OFs. Treatment with PDGFβ (50 ng/mL) for 48 hours robustly increased mRNA expression of *FOXC2*, *HMGA2*, and *HGF*, while significantly reducing *SFRP2* expression compared to vehicle controls (**Fig. 3A–D**). Cotreatment with sorafenib (250 nM) markedly blunted PDGFβ-induced *FOXC2*, *HMGA2*, and *HGF* expression, and restored *SFRP2* expression to baseline, indicating a broad inhibitory effect on PDGFβ-mediated transcriptional activation.

**Fig. 3.**
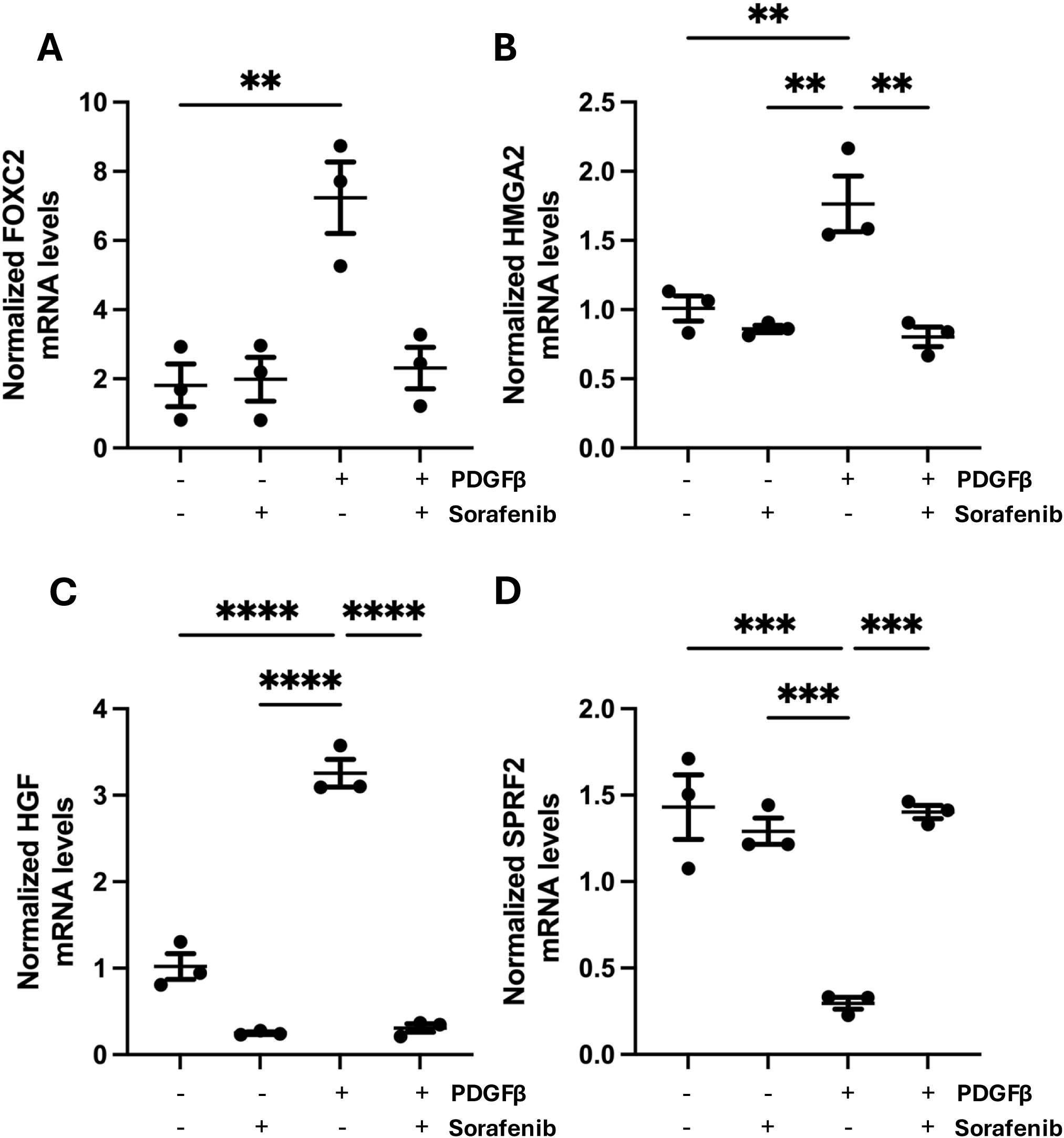
Sorafenib reverses PDGFβ-induced pathogenic gene expression in TED orbital fibroblasts. Orbital fibroblasts from two TED patient-derived strains were treated with vehicle, PDGFβ (50 ng/mL), or PDGFβ + sorafenib (250 nM) for 48 h. mRNA levels of *FOXC2* (A)*, HMGA2* (B)*, HGF* (C), and *SFRP2* (D) were measured by RT-qPCR and normalized to *18S rRNA* and *TBP*. PDGFβ significantly increased *FOXC2*, *HMGA2*, and *HGF* expression and reduced *SFRP2*; co-treatment with sorafenib blunted these effects. Bars represent mean ± SEM from two strains evaluated in triplicate (n = 6 per group). p < 0.05 (*), p < 0.001 (**), one-way ANOVA with post hoc tests.

To determine whether sorafenib’s inhibitory effects on gene expression were associated with attenuation of PDGFβ downstream signaling, we examined activation of AKT and thymidylate synthase expression by Western blot (28). PDGFβ stimulation increased phosphorylation of AKT, while total AKT levels remained unchanged (**Fig. 4A and B**). Sorafenib reduced PDGFβ-induced AKT phosphorylation and thymidylate synthase expression in a dose-dependent manner, with substantial inhibition observed at concentrations ≥125 nM (**Fig. 4B and C**). These findings suggest that sorafenib suppresses PDGFβ-mediated transcriptional responses, in part, by inhibiting AKT signaling downstream of PDGFβ.

**Fig. 4.**
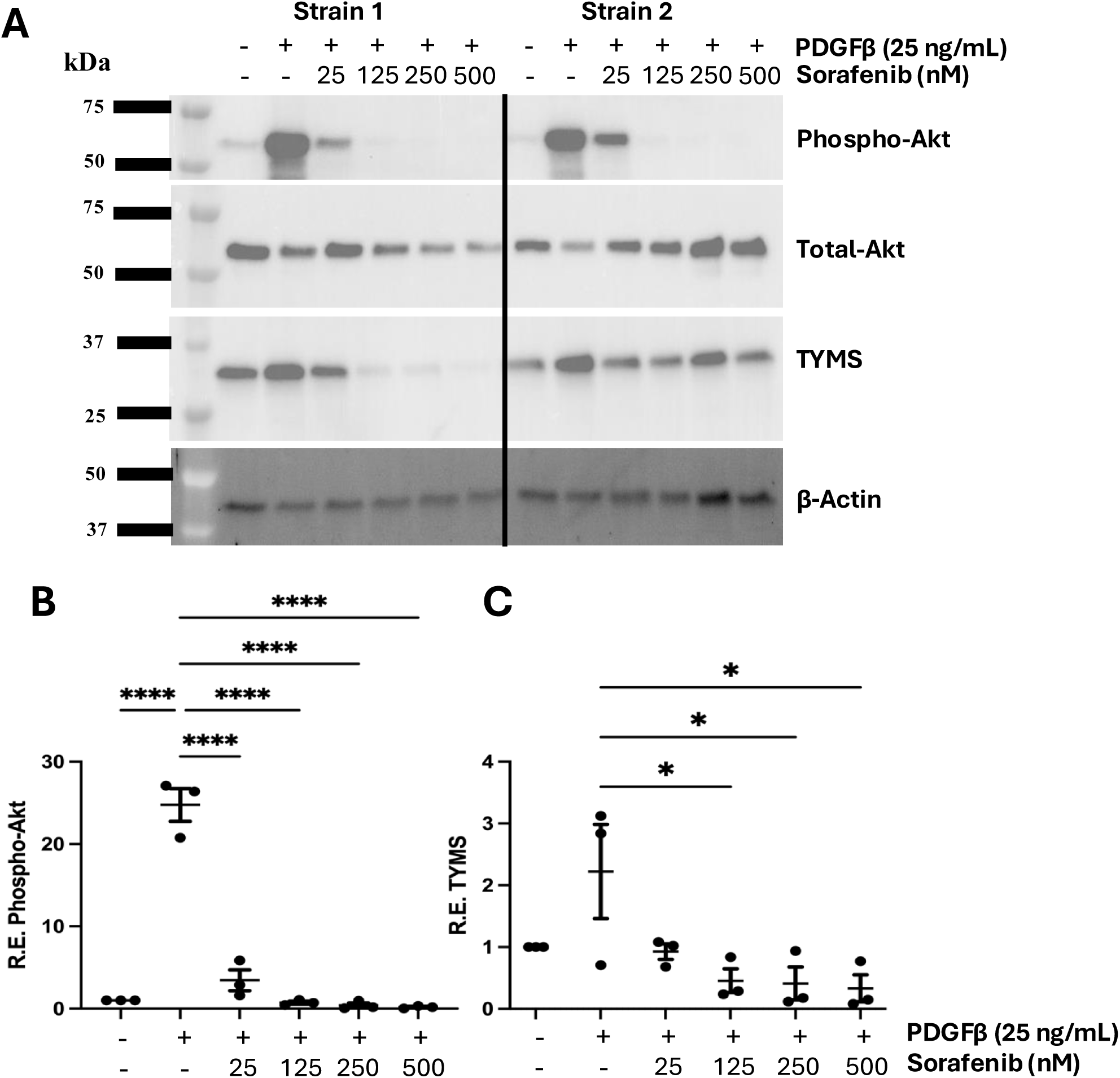
Sorafenib suppresses PDGFβ-induced activation of AKT and thymidylate synthase (TYMS) expression in TED orbital fibroblasts. (A) Representative Western blots of TED fibroblast strains treated with vehicle, PDGFβ (50 ng/mL), or PDGFβ plus increasing sorafenib concentrations (25–500 nM). PDGFβ robustly induced phosphorylation of AKT (phospho-AKT) and increased TYMS expression. Sorafenib reduced phospho-AKT and TYMS in a dose-dependent manner. Total AKT and β-Actin served as loading controls. (B) Quantification of relative phospho-AKT normalized to total AKT. (C) Quantification of TYMS normalized to β-Actin (Right panel). Data represent mean ± SEM from three independent TED strains (n = 3). p < 0.05 by one-way ANOVA.

### Sorafenib Blocks PDGFβ-Induced Hyaluronan Production in TED OFs

Excessive HA synthesis by OFs contributes to tissue expansion and edema in TED (16, 50, 51). Here, we tested PDGFβ’s ability to promote HA production in three different TED OF strains. After 48 hours of treatment, PDGFβ strongly induced HA production in all three TED OF strains evaluated, as measured by ELISA (**Supplemental Figure 5**). Sorafenib co-treatment suppressed HA levels in a dose-dependent manner, with maximal inhibition observed at 250 to 500 nM. Summary analysis across strains confirmed sorafenib’s consistent suppression of PDGFβ-induced HA production **(Fig. 5)**, indicating its potential to target a key pathogenic output of activated TED fibroblasts. Across three strains, the calculated IC₅₀ for sorafenib-mediated inhibition of PDGFβ-induced HA production was ∼100 nM, consistent with its inhibition of PDGFβ-AKT signaling.

**Fig. 5.**
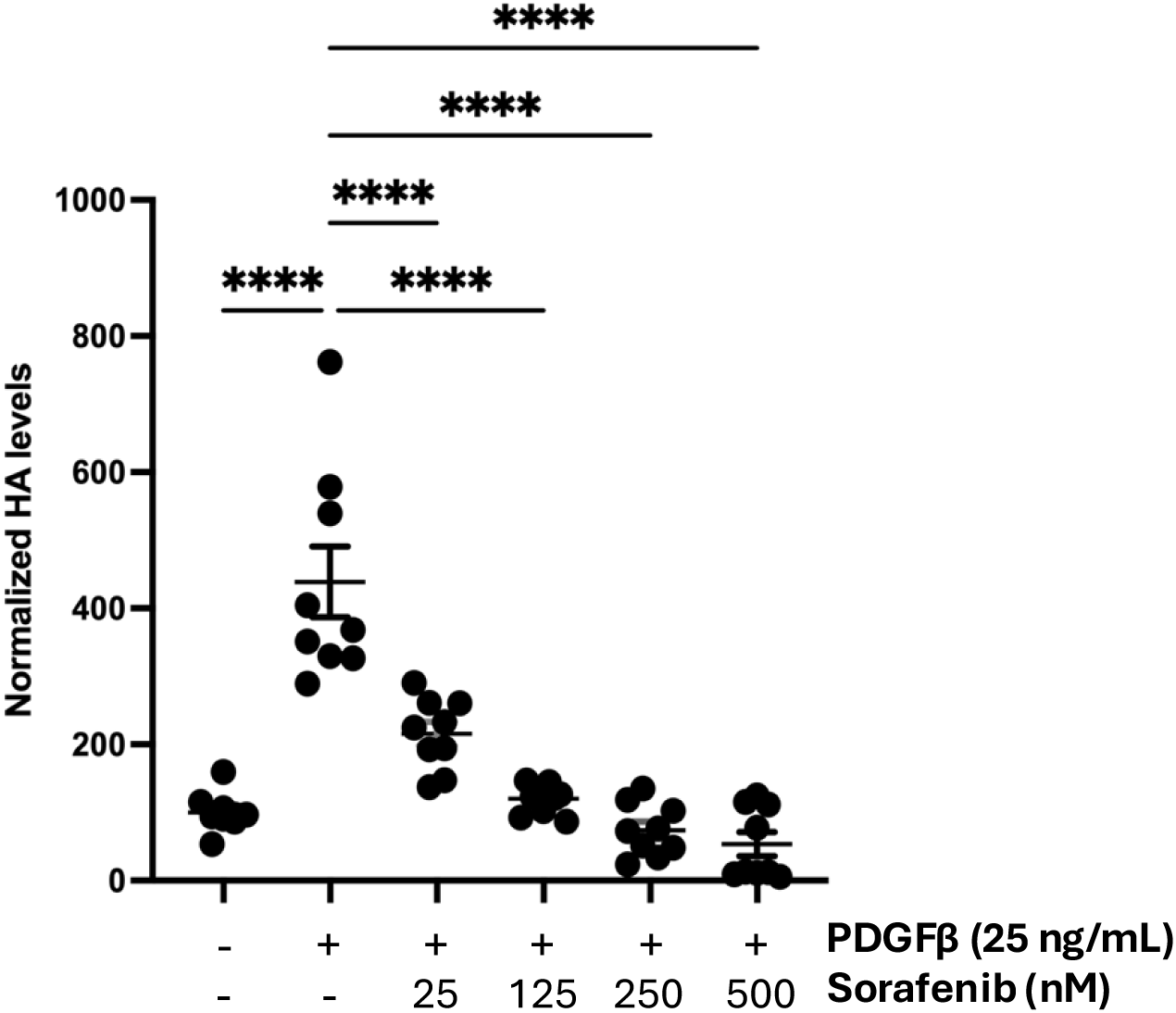
Sorafenib inhibits PDGFβ-induced hyaluronan (HA) production in TED orbital fibroblasts. TED fibroblasts from three independent patient-derived strains were treated with vehicle (Veh), PDGFβ (50 ng/mL), or PDGFβ (P) + sorafenib (25–500 nM) for 48 h. HA production was quantified. Individual strain responses show that PDGFβ markedly induces HA synthesis, which is suppressed by sorafenib in a dose-dependent manner. A summary graph combining normalized HA levels (Veh = 100%) across all strains demonstrates consistent inhibition at concentrations of 250 nM or greater. Data are mean ± SEM of triplicate wells from n = 3 strains.

## Discussion

This study provides an integrative transcriptomic and functional analysis directly comparing OFs from patients with TED and non-TED controls. Through unbiased RNA sequencing, we define a fibro-angiogenic phenotype in TED OFs characterized by coordinated upregulation of angiogenic and fibroblast activation programs. Among the most upregulated genes were *FOXC2*, *MET*, *HGF*, *HMGA2*, and *PLEKHG5*, which converge on PI3K/AKT signaling and angiogenic activation, while *SFRP2*, a negative regulator of Wnt signaling, was downregulated. This pattern is consistent with prior work demonstrating that PDGF receptors on stromal and perivascular cells drive proliferation and matrix remodeling through targetable signaling pathways (52). Together, these findings implicate the PI3K/AKT pathway as a convergence point that integrates multiple signaling cues to drive OF activation, proliferation, and HA synthesis. Notably, the prominence of angiogenic signatures challenges the established framing of TED as primarily an autoimmune-fibroinflammatory disorder, suggesting that fibrovascular crosstalk may be an underappreciated amplifier of orbital disease.

Building on prior work that identified IGF-1R and TSHR signaling as primary drivers of TED pathogenesis, our findings extend this paradigm by identifying PI3K/AKT as a key downstream node that integrates autoimmune (TSHR), growth factor (IGF-1R, PDGFR), and angiogenic (VEGFR, HGF/MET) responses to coordinate ECM remodeling and tissue expansion. The upregulation of SMAD3, BMP4, and TGFBR2 further links TGFβ/BMP signaling to this network, highlighting crosstalk between growth factor, angiogenic, and fibrotic pathways. This suggests a unified explanation for the diverse clinical manifestations of TED, including fibrosis, adipogenesis, and angiogenesis, within a shared PI3K/AKT-centered signaling network (**Fig. 6**).

**Fig. 6.**
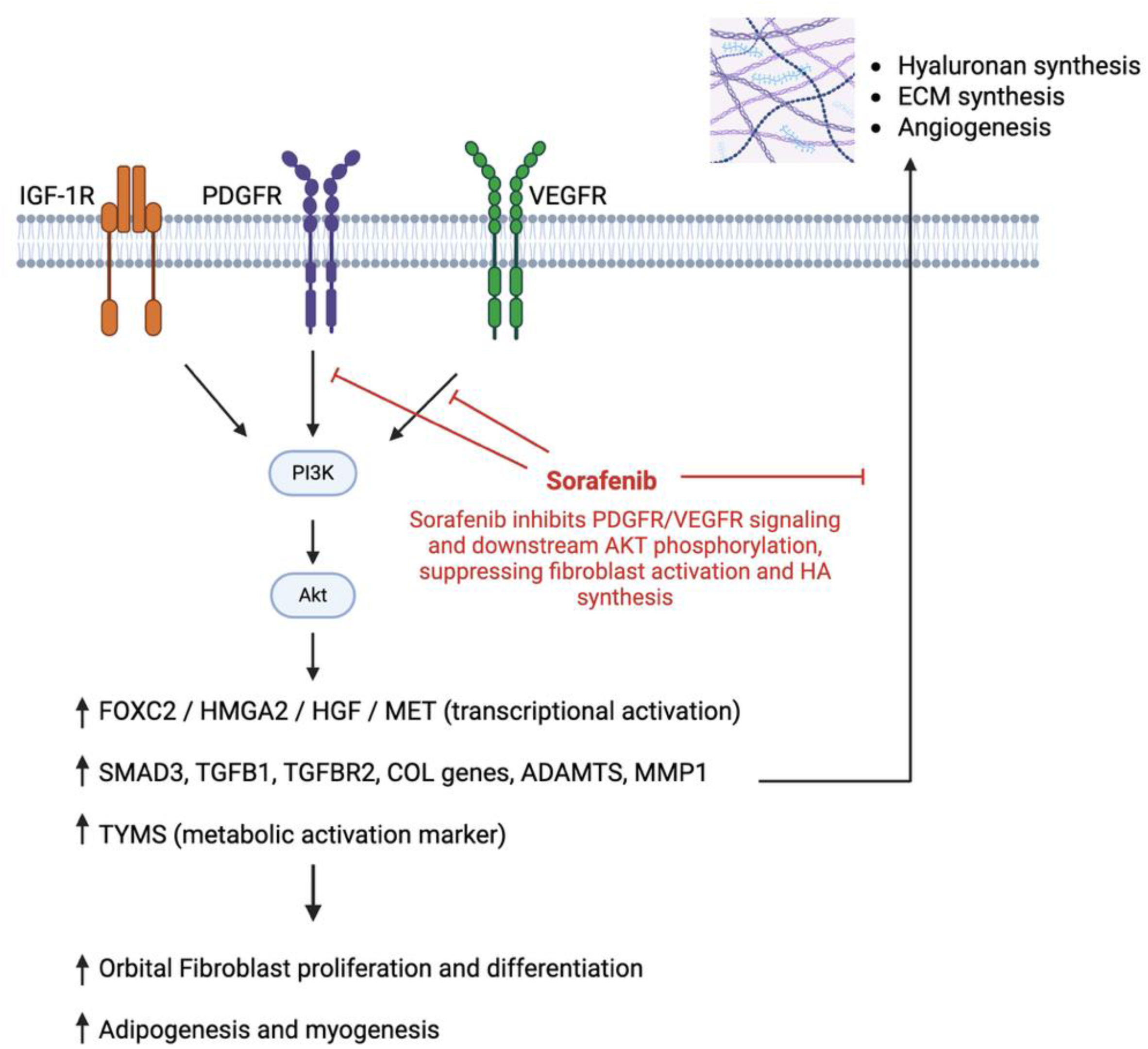
Proposed model illustrating PI3K/AKT and angiogenic pathway activation in TED orbital fibroblasts and sorafenib inhibition. Transcriptomic profiling revealed the activation of PI3K/AKT, ECM remodeling, and angiogenic pathways in TED OFs. PDGFR, VEGFR, and IGF-1R signaling converge on PI3K/AKT to promote the expression of FOXC2, HMGA2, HGF, and collagen-associated genes, thereby enhancing hyaluronan (HA) synthesis and tissue expansion. Sorafenib inhibits VEGFR/PDGFR signaling and downstream AKT activation, thereby suppressing fibroblast activation and extracellular matrix accumulation. Schematic created with BioRender.com.

Our Connectivity Map (LINCS L1000CDS²) analysis identified multiple FDA-approved kinase inhibitors predicted to reverse the TED gene expression profile, including mirdametinib, selumetinib, trametinib, and sorafenib. Functional studies confirmed that sorafenib, a multi-kinase inhibitor with established activity against Raf kinases, VEGFR, and PDGFRβ (53), dose-dependently suppressed PDGFβ-induced gene expression, AKT phosphorylation, and HA synthesis in TED OFs. Previous studies have revealed similar pathways in other systems; for example, in hepatic stellate cells, PDGFβ-driven activation and survival can be countered by sorafenib, in part by modulating Hippo/YAP signaling and inducing ferroptosis (48). Likewise, in both *in vitro* and *in vivo* settings, sorafenib blocked ossification of the posterior longitudinal ligament (OPLL) by disrupting vascularization and osteogenesis through VEGF/PDGF signaling pathways (49). These studies support the concept that PDGF-responsive, matrix-producing stromal cells are a druggable cell type for sorafenib, and our data extend this to OFs in TED.

Sorafenib also offers a therapeutic strategy that complements IGF-1R blockade with teprotumumab, potentially benefiting patients who are refractory to current treatments or relapse after therapy withdrawal. While teprotumumab primarily inhibits IGF-1R-mediated signaling to reduce inflammation and tissue expansion, sorafenib targets PI3K/AKT and angiogenic pathways, addressing additional drivers of fibroblast activation. This dual-pathway approach may enhance therapeutic efficacy and reduce reliance on long-term immunosuppression or invasive surgical interventions, such as orbital decompression, which carry significant risks and morbidity.

While our studies indicate that sorafenib may be effective in treating TED, this work serves as proof of principle for transcriptomic-guided drug repurposing in fibroinflammatory diseases. TED shares core pathogenic components with systemic sclerosis, idiopathic pulmonary fibrosis, and other organ-specific fibrotic diseases (persistent myofibroblast activation, ECM accumulation, growth factor dependence, and angiogenic crosstalk)(2). Sorafenib may also show therapeutic benefit in these diseases. Furthermore, the integrative strategy employed here: RNAseq ◊ pathway enrichment◊ LINCS analysis ◊ functional validation could be applied to uncover novel therapeutic targets for a broader spectrum of fibroblast-related disease.

This study also has limitations. The sample size was constrained by access to primary orbital tissue, and we employed bulk RNA-seq, which may mask fibroblast heterogeneity and immune-stromal interactions. Additional single-cell and spatial transcriptomic studies are needed to define TED-enriched fibroblast subsets and determine whether the angiogenic/ECM signature is confined to specific fibroblast subpopulations. *In vitro*, we focused on PDGFβ because it overlapped with the transcriptomic signal detected and with sorafenib’s target profile; however, other RTK ligands present in the orbit (IGF-1, VEGF, HGF/MET, and TGFβ family members) may also sustain the TED phenotype and should be further tested. Finally, sorafenib is a systemic multikinase inhibitor with known dermatologic, gastrointestinal, and metabolic toxicities in differentiated thyroid cancer (54); translating it to TED will require attention to dosing, schedule, and possibly local/topical or depot delivery to minimize off-target effects. Alternatively, more selective multikinase/PI3K inhibitors with reduced adverse profiles may emerge as preferred agents.

In summary, we redefine TED OFs as PI3K/AKT- and angiogenesis-driven orchestrators of TED pathology rather than solely inflammatory/fibrotic targets and demonstrate that an FDA-approved multi-kinase inhibitor can pharmacologically reverse the TED transcriptional state. The alignment of RNAseq findings, pathway predictions, and functional validation establishes sorafenib as a plausible candidate for clinical evaluation in TED, particularly for patients with incomplete responses or relapse after IGF-1R blockade. More broadly, this integrated discovery-to-validation pipeline illustrates how transcriptomic profiling can be translated into mechanism-based therapy design across fibroblast-driven disorders.

## Methods

*Sex as a biological variable. Patient demographics, including sex and age, are detailed in* ***Table 1****. Due to the limited sample size, sex-based comparisons of human orbital fibroblast (OF) transcriptomic data were not performed*.

**Table 1:** Demographic Information for non-TED vs. TED Subjects.

| Category | Non-TED | TED |
| --- | --- | --- |
| Age (Mean) range (yrs) | (52.5) 45–58 | (54.3) 27-69 |
| Smokers (N) | 1 | 4 |
| Steroid Use (N) | 0 | 3 |
| Sex (M, F) | 1,3 | 2,5 |
| Total | 4 | 7 |

### Tissue Procurement and Study Approval

*OFs* were isolated from orbital fat specimens obtained during decompression surgery from patients with thyroid eye disease (TED) (n = 7) or from patients undergoing orbital surgery for non-TED indications (n = 4). All human tissue collection procedures were conducted in accordance with the Declaration of Helsinki and approved by the University of Rochester Institutional Review Board. Written informed consent was obtained from all participants. Donor demographics are summarized in **Table 1**. Primary cultures were established from fresh tissue using explant outgrowth. Tissue fragments (∼1 mm³) were placed in 35-mm dishes containing Dulbecco’s Modified Eagle Medium/Nutrient Mixture F-12 (DMEM/F12; Gibco) supplemented with 10% fetal bovine serum (FBS; Sigma), 2 mM L-glutamine, and antibiotic-antimycotic (100 U/mL penicillin, 100 μg/mL streptomycin, 0.25 μg/mL amphotericin B; Gibco). Cultures were maintained at 37 °C in a humidified 5% CO₂ incubator. Fibroblasts migrating from explants were expanded to confluence (approximately 2–3 weeks), tissue remnants were removed, and adherent cells were passaged using 0.05% trypsin-EDTA (Gibco). Cell identity was verified by vimentin⁺/cytokeratin⁻/CD45⁻. Experiments used cells between passages 4 and 10.

### RNA Isolation and Sequencing

For RNA-seq, total RNA was extracted using the RNeasy Mini Kit (Qiagen) with on-column DNase digestion (RNase-Free DNase Set, Qiagen). RNA quantity and purity were assessed by NanoDrop spectrophotometry, and integrity by Agilent 2100 Bioanalyzer; only samples with RIN > 8.0 were used. Libraries were prepared with the TruSeq Stranded mRNA Library Prep Kit (Illumina) at the University of Rochester Functional Genomics Core and sequenced on an Illumina NovaSeq 6000 (150-bp paired-end; ∼30 million reads/sample).

### Bioinformatics and Differential Expression Analysis

Reads were processed and analyzed with DESeq2 (R). Differential expression between TED and non-TED OFs was assessed with Benjamini–Hochberg correction; adjusted p < 0.05 was considered significant. Gene ontology (GO) and pathway enrichment were performed using Enrichr, ToppGene, and DAVID. KEGG and Reactome were the primary pathway databases queried (adjusted p < 0.05).

### LINCS Analysis

Differentially expressed genes were queried in the Library of Integrated Network-Based Cellular Signatures (LINCS) database. The L1000 characteristic direction signature search engine (CDS²) platform was used to identify compounds predicted to reverse the transcriptional signature found in TED OFs. Top candidates were selected based on their relevance to TED-associated pathways.

### PDGFβ and Sorafenib Treatment of Orbital Fibroblasts

For stimulation experiments, OFs at ∼70% confluence were washed twice with PBS and placed in DMEM/F12 containing 0.1% FBS with 50 ng/mL recombinant human PDGF-BB (hereafter referred to as PDGFβ) (PeproTech, Cranbury, NJ, USA) for 48 hours at 37 °C, 5% CO_2_. Vehicle controls received the corresponding PDGFβ diluent (0.1% BSA in PBS). Conditioned media were collected for HA ELISA, and cells were harvested for Western blotting and RNA isolation as indicated below. Sorafenib (BAY-43-9006; MedChemExpress) was dissolved in DMSO to prepare stock solutions, which were diluted into 0.1% FBS DMEM/F12. Cells were pre-treated with sorafenib (25–500 nM) for 30 minutes, after which PDGFβ was added without media change; sorafenib was maintained for the duration of the PDGFβ exposure (48 hr unless otherwise indicated). The final DMSO concentration was ≤0.1% (v/v). Vehicle controls received DMSO at the matched final concentration.

### Hyaluronic acid (HA) ELISA

HA in conditioned media was quantified using the HA ELISA kit (Echelon Biosciences, Cat. K-1200) according to the manufacturer’s instructions. The media were centrifuged at 300 × g for 5 min to remove debris and then diluted 1:5 in assay buffer. A standard curve (1 MDa HA; 0–5,000 ng/mL) was run in parallel, and samples were assayed in triplicate. Briefly, samples or standards were first incubated with the kit HA detector (biotinylated HA-binding protein) to allow competitive binding of HA. The mixtures were then transferred to the supplied HA ELISA plate, incubated, and washed as instructed. They were then incubated for 30 min with the kit’s streptavidin–alkaline phosphatase reagent. After three washes, plates were developed with 100 μL p-nitrophenyl phosphate (PNPP) for 25 min. Reactions were stopped, and absorbance was read at 405 nm (reference 610 nm) on a Bio-Rad iMark microplate reader. HA concentrations were interpolated from the standard curve.

### Western Blotting

Cells were washed with ice-cold PBS and then lysed in 50 mM Tris-HCl (pH 6.8), 2% SDS, supplemented with protease and phosphatase inhibitor cocktails (Cell Signaling Technology). Lysates were heated at 95 °C for 5 min, sonicated briefly, and then clarified (14,000 × g, 10 min, 4 °C). Protein concentration was determined using the DC Protein Assay (Bio-Rad). Equal amounts (5–10 µg) were separated on 4– 20% Mini-PROTEAN TGX Stain-Free gels (Bio-Rad) and transferred to 0.45 µm Immobilon-PVDF (Millipore). Membranes were blocked in 5% non-fat dry milk/0.1% Tween-20 in TBS (TBS-T) for 1 h and incubated overnight (4 °C) with primary antibodies: thymidylate synthase (TYMS; rabbit, CST #3766, 1:1,000), phospho-AKT (Ser473; rabbit, CST #4060, 1:1,000), total AKT (rabbit, CST #9272, 1:1,000), and β-tubulin (anti-tubulin-hFAB Rhodamine, Bio-Rad #12004165, 1:5,000). After TBS-T washes, membranes probed for TYMS, pAKT, and AKT were incubated for 1 h with HRP-conjugated anti-rabbit secondary antibody (Jackson ImmunoResearch, 1:5,000), developed with Immobilon Western Chemiluminescent HRP Substrate (Millipore), and imaged on a ChemiDoc MP system (Bio-Rad). β-tubulin was detected by direct fluorescence (rhodamine channel) on the same system. Band intensities were quantified with Image Lab software (Bio-Rad). Equal loading was verified by tubulin and stain-free total protein imaging.

### Real-Time Quantitative PCR (RT-qPCR)

Total RNA was extracted with QIAzol Lysis Reagent and purified using the RNeasy Mini Kit (Qiagen) according to the manufacturer’s protocol. RNA concentration and purity (A260/A280) were measured on a DS-11+ spectrophotometer (DeNovix). cDNA was synthesized from 500 ng RNA using iScript™ Reverse Transcription Supermix (Bio-Rad). qPCR was performed on a CFX Connect™ Real-Time PCR Detection System (Bio-Rad) using SsoFast™ EvaGreen Supermix (Bio-Rad) and gene-specific primers. Target genes were *FOXC2, HGF, HMGA2, SFRP2, PLEKHG5, and MET*. Relative mRNA expression was calculated by the 2^-ΔΔCq^ method, normalized to the geometric mean of *18S rRNA* and *TBP* Cq values. Primer sequences are provided in **Table 2**.

**Table 2:**
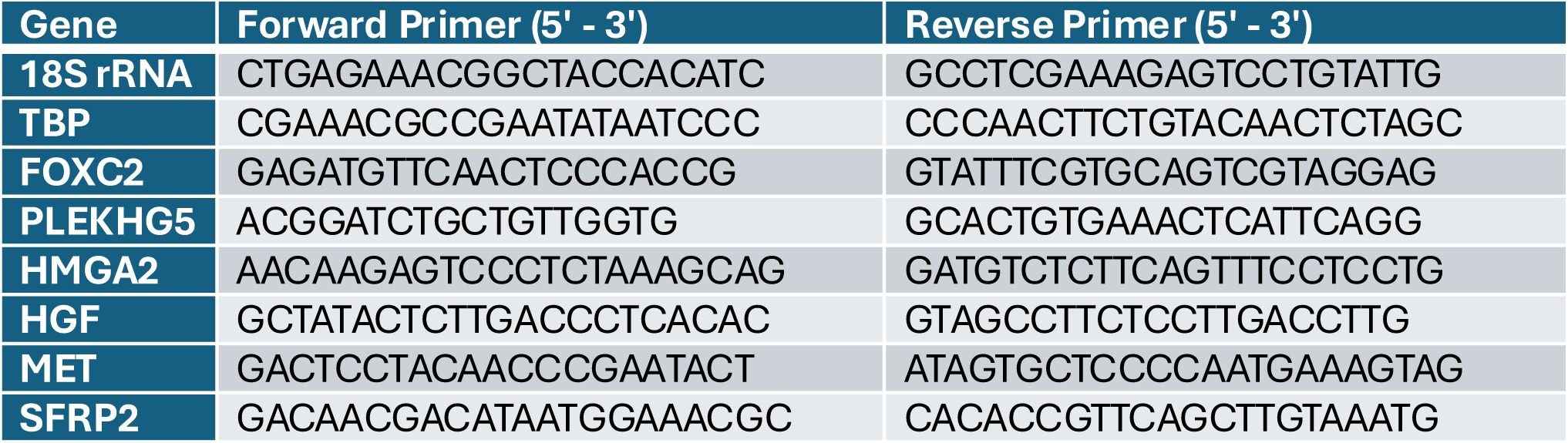
Gene primer sequences for RT-qPCR.

**Table 3.** FDA-approved drugs predicted to reverse the TED transcriptional signature by LINCS L1000CDS² analysis. Rank indicates the position of each compound among all screened agents. The cosine distance reflects the inverse correlation between each drug’s gene expression signature and the TED versus non-TED differential expression profile; higher values indicate a more substantial predicted reversal of the signature. Selected top-ranked compounds are shown.

| Rank | Cosine Distance | Drug | Mechanism |
| --- | --- | --- | --- |
| 1 | 1.4617 | Mirdametinib | MEK1/2 inhibitor |
| 2 | 1.4521 | Selumetinib | MEK1/2 inhibitor |
| 5 | 1.4346 | Trametinib | MEK1/2 inhibitor |
| 20 | 1.3818 | Sorafenib | Multikinase inhibitor (Raf, RET, VEGFR, PDGFR) |
| 22 | 1.3805 | Hydrocortisone Valerate | Topical glucocorticoid |
| 24 | 1.3778 | Beclomethasone Dipropionate | Inhaled/Topical glucocorticoid |
| 28 | 1.3721 | Afatinib | ErbB family inhibitor (EGFR, HER2, HER4) |
| 34 | 1.3625 | Quinidine Gluconate | Antiarrhythmic |

### Statistics

HA ELISA, Western blot densitometry, and RT-qPCR data were analyzed in GraphPad Prism (v10). A two-tailed unpaired Student’s *t-test* was used for comparisons between two groups. For comparisons with ≥3 groups, one-way ANOVA followed by Tukey’s post hoc test was used; p < 0.05 was considered significant. Sorafenib IC_50_ values were calculated from normalized HA levels using a four-parameter logistic (4PL) fit in Prism. All *in vitro* experiments were performed in at least two independent TED orbital fibroblast strains, with technical triplicates per condition. Data are presented as mean ± SEM.

## Study approval

This study was performed in accordance with protocols (URMC00099) approved by the IRB at the University of Rochester Medical Center. All participants provided written informed consent.

## Data availability

The raw and processed RNA-sequencing data generated in this study have been deposited in the NCBI Gene Expression Omnibus (GEO). The final GEO accession number (GSE) will be provided upon assignment. Supporting data for all graphs and blots presented in the figures are available in the Supporting Data file. Supplemental Figures 1-4: Uncropped Western blot images for phospho-AKT, AKT, TYMS, and β-Actin. Supplemental Figure 5: Individual Strains HA ELISA. Supplemental Figure 6: Clustering and PCA analysis of TED vs Non-TED OF samples. Supplemental Table 1: Complete KEGG (p-adj < 0.05) and Reactome (p-adj < 0.10) enrichment plots for TED DEGs.

## Statements and Declarations

### Author Contributions

Material preparation, data collection, and analysis were performed by Kyle Yuan, Emma Ushchak, Phillip Truong, Charkira C. Patrick, Elisa Roztocil, Steven E. Feldon, and Collynn F. Woeller. Kyle Yuan wrote the first draft of the manuscript. All authors read and approved the final manuscript.

### Ethics Approval

This study was performed in line with the principles of the Declaration of Helsinki. Approval was granted by the University of Rochester IRB, Protocol Number STUDY00000099.

### Consent to Participate

Informed consent was obtained from all individual participants included in the study.

