## Supplemental for "Transcriptomic Profiling of Thyroid Eye Disease Orbital Fibroblasts Identifies Sorafenib as a Novel Therapeutic"

### Supplemental Data

Supplemental Figure 1

Sorafenib Dose-Response (-/+ PDGF) - S080 / S084 TED OF –
**phospho-Akt** (uncropped blot)

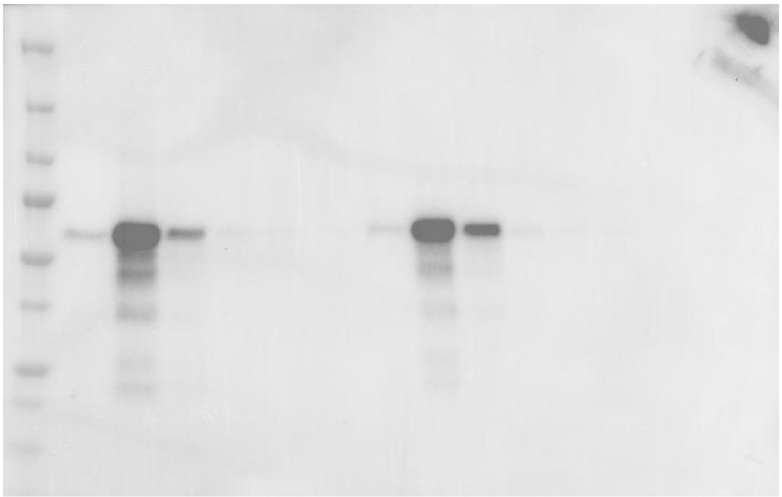

Sorafenib Dose-Response (-/+ PDGF) - S080 / S084 TED OF –
**total-Akt** (uncropped blot)

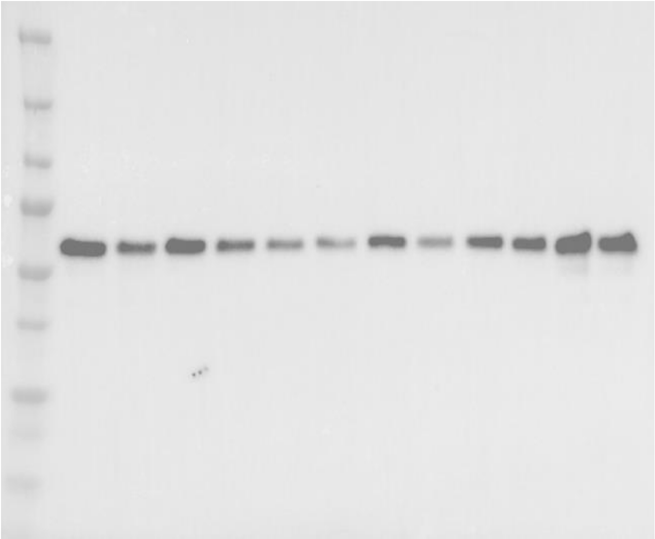

Sorafenib Dose-Response (-/+ PDGF) - S080 / S084 TED OF –
**TYMS** (uncropped blot)

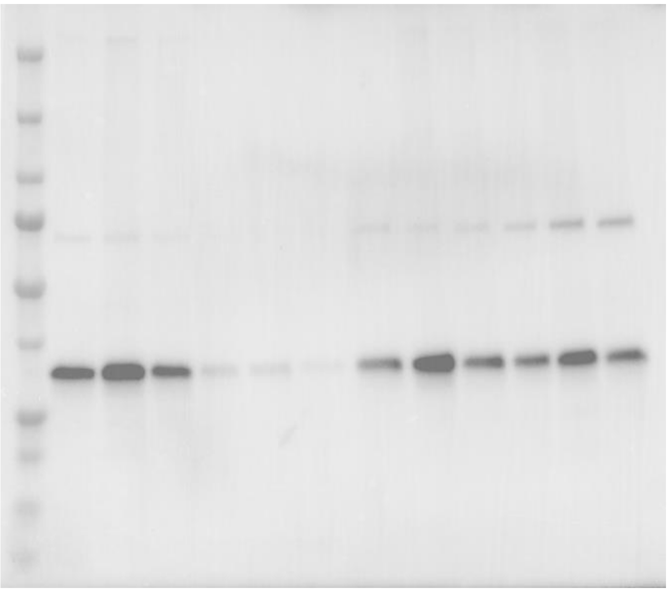

Supplemental Figure 4

Sorafenib Dose-Response (-/+ PDGF) - S080 / S084 TED OF – **<u>β-Actin</u>** (uncropped blot)

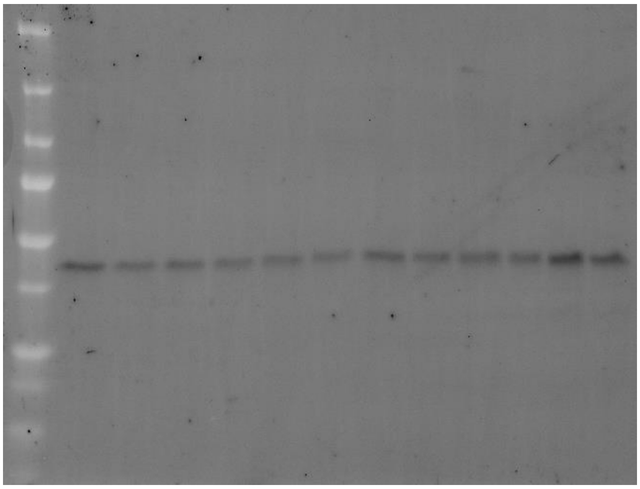

Supplemental Figure 5

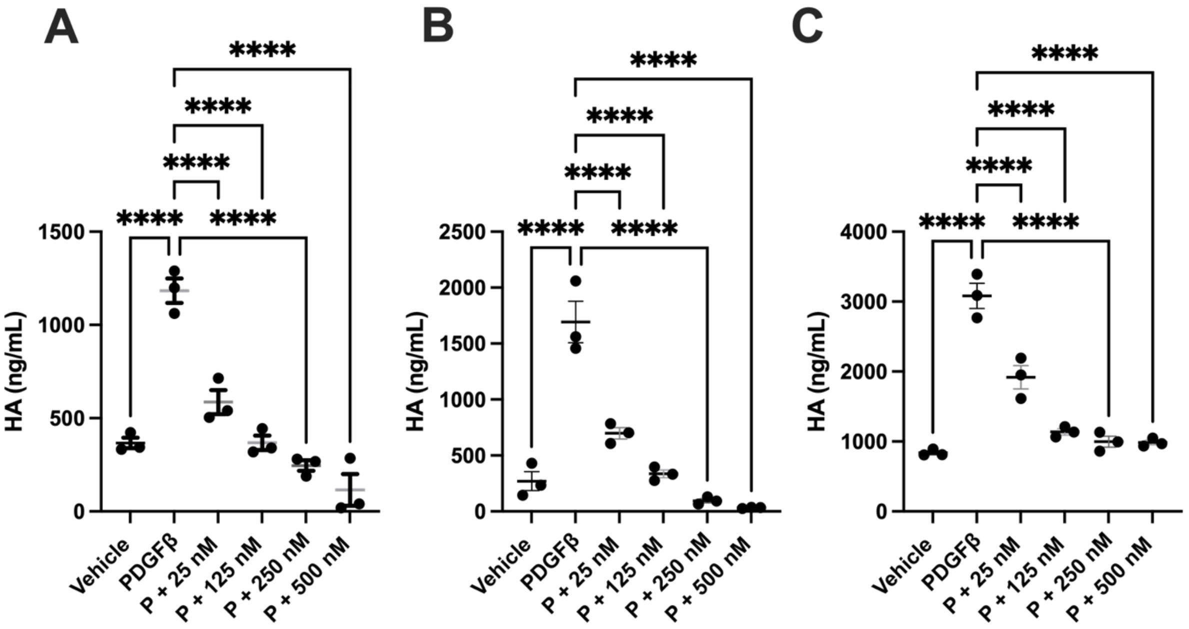

### Supplemental Figure 6

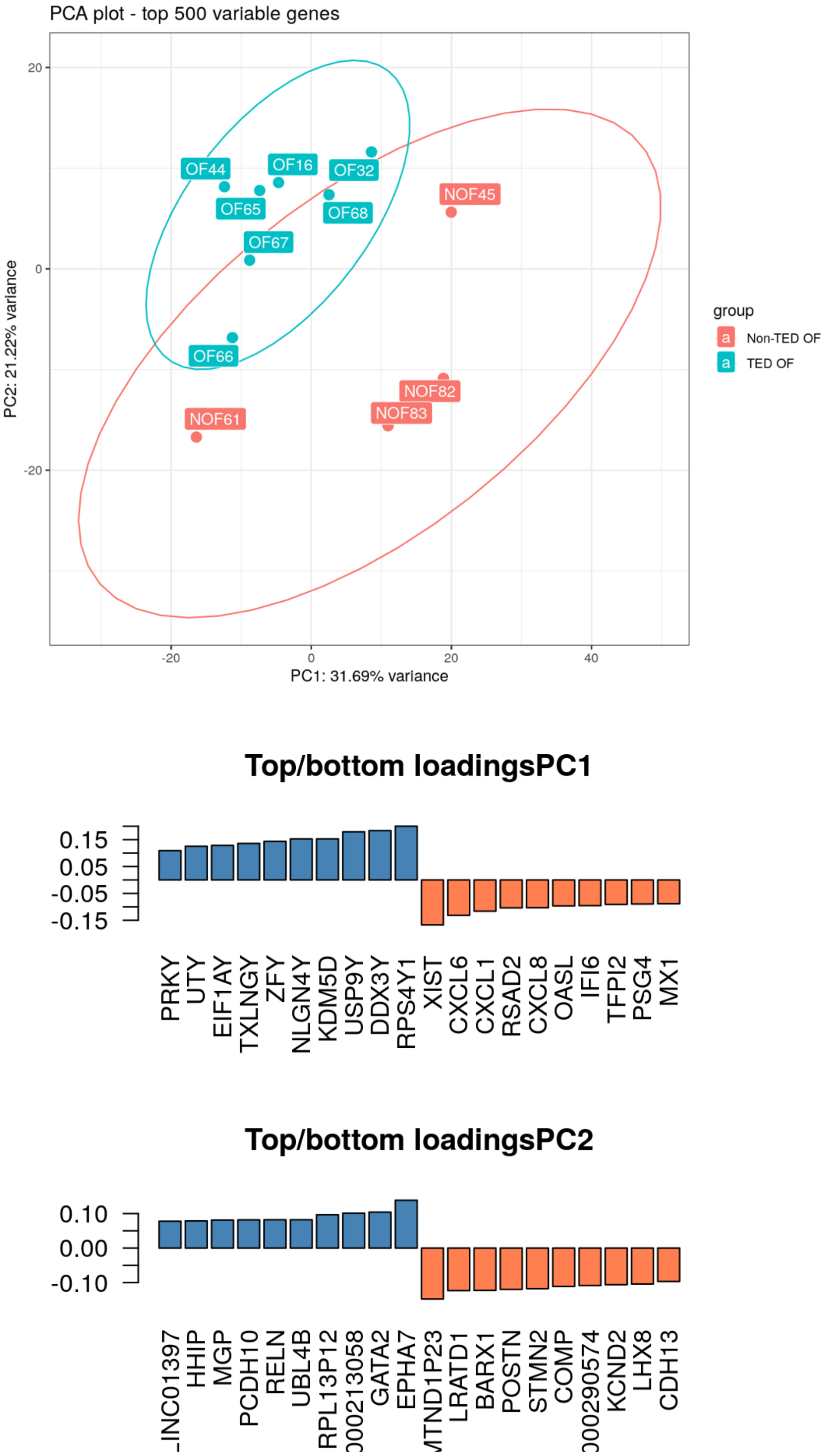

### Supplemental Table 1

#### KEGG p-adj 0.05

| Term | P-value | Adjusted p-value | Odds Ratio | Combined Score | Overlap | Genes |
|---|---|---|---|---|---|---|
| Pathways in cancer | 1.18E-06 | 0.000255266 | 3.740260821 | 51.06616007 | 21/531 | LAMA5, JUN, CAMK2D, SMAD3, TGFB1, JUP, MMP1, HHIP, HGF, PLEKHG5, PIK3CB, TRAF1, TGFBR2, BMP4, FGF16, SMO, SUFU, COL4A6, RARB, COL4A5, MET |
| TGF-beta signaling pathway | 1.42E-05 | 0.00153568 | 8.136847545 | 90.85223342 | 8/94 | BMP4, SMAD3, TGFB1, PPP2R1A, GDF6, GDF5, TGFBR2, GDF7 |
| Gastric cancer | 6.43E-05 | 0.004651716 | 5.632748724 | 54.36618769 | 9/149 | FGF16, SMAD3, TGFB1, JUP, HGF, RARB, PIK3CB, MET, TGFBR2 |
| Rheumatoid arthritis | 0.000107749 | 0.005604077 | 7.088238321 | 64.75603465 | 7/93 | IL11, JUN, ATP6V1G2, TGFB1, MMP1, TNFSF11, TLR2 |
| Focal adhesion | 0.000129126 | 0.005604077 | 4.596060386 | 41.15643579 | 10/201 | LAMA5, JUN, RELN, HGF, CAPN2, COL4A6, COL4A5, COL9A3, PIK3CB, MET |
| AGE-RAGE signaling pathway in diabetic complications | 0.000169963 | 0.006146989 | 6.552383671 | 56.87423677 | 7/100 | JUN, SMAD3, TGFB1, COL4A6, COL4A5, PIK3CB, TGFBR2 |
| Hippo signaling pathway | 0.000664901 | 0.018391849 | 4.498810036 | 32.91271992 | 8/163 | BMP4, SMAD3, TGFB1, PPP2R1A, GDF6, GDF5, GDF7, TGFBR2 |
| Small cell lung cancer | 0.000725563 | 0.018391849 | 6.048867944 | 43.72462188 | 6/92 | LAMA5, COL4A6, COL4A5, RARB, PIK3CB, TRAF1 |
| Relaxin signaling pathway | 0.000798719 | 0.018391849 | 4.987487306 | 35.5732586 | 7/129 | JUN, TGFB1, MMP1, COL4A6, COL4A5, PIK3CB, TGFBR2 |
| PI3K-Akt signaling pathway | 0.000958063 | 0.018391849 | 3.084067635 | 21.43611014 | 12/354 | FGF16, LAMA5, RELN, PPP2R1A, HGF, COL4A6, COL4A5, THEM4, COL9A3, PIK3CB, MET, TLR2 |
| Amoebiasis | 0.001244501 | 0.018391849 | 5.416024229 | 36.22789641 | 6/102 | LAMA5, TGFB1, COL4A6, COL4A5, PIK3CB, TLR2 |
| Chagas disease | 0.001244501 | 0.018391849 | 5.416024229 | 36.22789641 | 6/102 | JUN, TGFB1, PPP2R1A, PIK3CB, TGFBR2, TLR2 |
| Amphetamine addiction | 0.001260235 | 0.018391849 | 6.751302083 | 45.07477846 | 5/69 | CAMK2D, GRIN2A, JUN, FOSB, STX1A |
| Renal cell carcinoma | 0.001260235 | 0.018391849 | 6.751302083 | 45.07477846 | 5/69 | JUN, TGFB1, HGF, PIK3CB, MET |
| Protein digestion and absorption | 0.001308846 | 0.018391849 | 5.359916436 | 35.58239202 | 6/103 | COL13A1, COL25A1, COL4A6, COL4A5, COL9A3, PRSS3 |
| Axon guidance | 0.001356081 | 0.018391849 | 4.003678161 | 26.43691287 | 8/182 | SEMA5A, EPHA7, CAMK2D, SMO, LRRC4, PIK3CB, MET, GDF7 |
| Transcriptional misregulation in cancer | 0.001900435 | 0.024258491 | 3.784154589 | 23.71027367 | 8/192 | HHEX, JUP, LMO2, HMGA2, PLAT, TRAF1, MET, TGFBR2 |
| Calcium signaling pathway | 0.002078174 | 0.025053545 | 3.397959184 | 20.98669821 | 9/240 | FGF16, GRIN2A, HRH1, CAMK2D, MCOLN3, HGF, ADRA1B, MET, MCOLN2 |
| Malaria | 0.002716974 | 0.031030707 | 7.488513385 | 44.24390729 | 4/50 | TGFB1, HGF, MET, TLR2 |
| Hepatitis B | 0.002951053 | 0.032018922 | 3.919040822 | 22.830738 | 7/162 | JUN, EGR3, SMAD3, TGFB1, PIK3CB, TGFBR2, TLR2 |
| Colorectal cancer | 0.003330052 | 0.034410536 | 5.329759584 | 30.40503873 | 5/86 | JUN, SMAD3, TGFB1, PIK3CB, TGFBR2 |
| ECM-receptor interaction | 0.003676562 | 0.035473351 | 5.200803213 | 29.15454459 | 5/88 | LAMA5, RELN, COL4A6, COL4A5, COL9A3 |
| Osteoclast differentiation | 0.003759848 | 0.035473351 | 4.29154986 | 23.96133936 | 6/127 | JUN, TGFB1, TNFSF11, FOSB, PIK3CB, TGFBR2 |

#### Reactome p adj 0.1

| Term | P-value | Adjusted P-value | Odds Ratio | Combined Score | Overlap | Genes |
|---|---|---|---|---|---|---|
| Extracellular Matrix Organization | 0.000000505 | 0.000385271 | 5.058220289 | 73.33823686 | 16/300 | LAMA5, TGFB1, COL13A1, PCOLCE2, COL25A1, MMP1, ICAM5, LTBP3, GDF5, BMP4, CAPN2, COL4A6, COL4A5, ADAM9, COL9A3, ADAMTS8 |
| Degradation of the Extracellular Matrix | 0.00000581 | 0.002215807 | 6.773715074 | 81.66559862 | 10/140 | LAMA5, COL13A1, COL25A1, MMP1, CAPN2, COL4A6, ADAM9, COL4A5, COL9A3, ADAMTS8 |
| Collagen Degradation | 0.0000094 | 0.002391143 | 10.71029343 | 123.96768 | 7/64 | COL13A1, COL25A1, MMP1, COL4A6, ADAM9, COL4A5, COL9A3 |
| Signaling by Receptor Tyrosine Kinases | 0.0000182 | 0.003464654 | 3.305826208 | 36.0867498 | 19/536 | CYFIP2, LAMA5, ATP6V1G2, EGR3, JUP, HGF, PLAT, PIK3CB, SHB, FGF16, PPP2R1A, POLR2E, FOSB, COL4A5, THEM4, COL9A3, TRIB1, MET, PTPN3 |
| SMAD2 3 Phosphorylation Motif Mutants in Cancer | 0.0000304 | 0.003482629 | 85.93043478 | 893.712273 | 3/6 | SMAD3, TGFB1, TGFBR2 |
| TGFBR1 KD Mutants in Cancer | 0.0000304 | 0.003482629 | 85.93043478 | 893.712273 | 3/6 | SMAD3, TGFB1, TGFBR2 |
| Signal Transduction | 0.000032 | 0.003482629 | 1.979103733 | 20.48632772 | 53/2613 | CYFIP2, HHIP, PLAT, MSI2, PIK3CB, SHB, ADRA1B, GJA1, BAIAP2L1, PPP2R1A, SUFU, ARHGDIB, PIP4K2C, PLEKHG3, DUSP5, ATP6V1G2, HGF, PLEKHG5, TRAF1, TGFBR2, SMO, LINC01139, RARB, COL4A5, TRIB1, MET, GNAZ, LAMA5, RGS17, CAMK2D, CRABP2, LTBP3, FAM169A, ARHGAP22, NEURL1B, HRH1, P2RY6, HCAR1, POLR2E, THEM4, GPC5, JUN, EGR3, SMAD3, TGFB1, JUP, FGF16, PDE10A, DHRS9, FOSB, COL9A3, RGS12, PTPN3 |
| Loss of Function of SMAD2 3 in Cancer | 0.0000528 | 0.004474274 | 64.44456522 | 634.7431775 | 3/7 | SMAD3, TGFB1, TGFBR2 |
| Loss of Function of TGFBR1 in Cancer | 0.0000528 | 0.004474274 | 64.44456522 | 634.7431775 | 3/7 | SMAD3, TGFB1, TGFBR2 |
| Signaling by TGF-beta Receptor Complex in Cancer | 0.0000837 | 0.006387523 | 51.55304348 | 483.9841857 | 3/8 | SMAD3, TGFB1, TGFBR2 |
| Collagen Biosynthesis and Modifying Enzymes | 0.000129255 | 0.008965612 | 8.538744854 | 76.45354477 | 6/67 | COL13A1, PCOLCE2, COL25A1, COL4A6, COL4A5, COL9A3 |
| Collagen Chain Trimerization | 0.000154143 | 0.009800909 | 11.09311741 | 97.37129714 | 5/44 | COL13A1, COL25A1, COL4A6, COL4A5, COL9A3 |
| Negative Regulation of the PI3K AKT Network | 0.000469198 | 0.027538334 | 5.484812246 | 42.038261 | 7/118 | FGF16, PPP2R1A, HGF, THEM4, PIP4K2C, PIK3CB, MET |
| Collagen Formation | 0.000645844 | 0.035198493 | 6.193517936 | 45.49109632 | 6/90 | COL13A1, PCOLCE2, COL25A1, COL4A6, COL4A5, COL9A3 |
| Molecules Associated With Elastic Fibres | 0.000876718 | 0.041808503 | 10.44541485 | 73.52866821 | 4/37 | BMP4, TGFB1, LTBP3, GDF5 |
| HSF1-dependent Transactivation | 0.000876718 | 0.041808503 | 10.44541485 | 73.52866821 | 4/37 | CAMK2D, HSPA6, COL4A6, TNFRSF21 |
| Germ Layer Formation at Gastrulation | 0.000940677 | 0.042219775 | 18.40341615 | 128.2517738 | 3/17 | EOMES, BMP4, SMAD3 |
| MET Activates PTPN11 | 0.001320486 | 0.053027926 | 57.03896104 | 378.1543745 | 2/5 | HGF, MET |
| MET Interacts With TNS Proteins | 0.001320486 | 0.053027926 | 57.03896104 | 378.1543745 | 2/5 | HGF, MET |
| Elastic Fibre Formation | 0.001691013 | 0.060924483 | 8.61441048 | 54.98085013 | 4/44 | BMP4, TGFB1, LTBP3, GDF5 |
| PI5P, PP2A and IER3 Regulate PI3K AKT Signaling | 0.001919451 | 0.060924483 | 4.949528005 | 30.9628411 | 6/111 | FGF16, PPP2R1A, HGF, PIP4K2C, PIK3CB, MET |
| ECM Proteoglycans | 0.001939946 | 0.060924483 | 6.083518656 | 37.99215296 | 5/76 | LAMA5, TGFB1, COL4A6, COL4A5, COL9A3 |
| MET Receptor Activation | 0.001965523 | 0.060924483 | 42.77705628 | 266.5864874 | 2/6 | HGF, MET |
| MET Activates PI3K AKT Signaling | 0.001965523 | 0.060924483 | 42.77705628 | 266.5864874 | 2/6 | HGF, MET |
| Kidney Development | 0.001996215 | 0.060924483 | 8.203368684 | 50.99626094 | 4/46 | IRX1, BMP4, FOXC2, IRX2 |
| TGF-beta Receptor Signaling Activates SMADs | 0.002162223 | 0.063452942 | 8.012186453 | 49.16772936 | 4/47 | SMAD3, TGFB1, LTBP3, TGFBR2 |
| Long-term Potentiation | 0.002326925 | 0.065757175 | 12.87847826 | 78.08488791 | 3/23 | CAMK2D, GRIN2A, NRGN |
| RUNX3 Regulates CDKN1A Transcription | 0.002730626 | 0.07440955 | 34.21991342 | 202.0078317 | 2/7 | SMAD3, TGFB1 |
| PI Metabolism | 0.003007686 | 0.079133244 | 5.465245392 | 31.73440875 | 5/84 | PIK3CB, PIP4K2C, PLEKHA4, SYNJ2, ENPP6 |
| Integrin Cell Surface Interactions | 0.003165912 | 0.08016573 | 5.396655702 | 31.05944818 | 5/85 | COL13A1, COL4A6, COL4A5, COL9A3, ICAM5 |
| Downregulation of TGF-beta Receptor Signaling | 0.003329521 | 0.08016573 | 11.19697543 | 63.87792635 | 3/26 | SMAD3, TGFB1, TGFBR2 |
| Synthesis of PIPs at the Plasma Membrane | 0.003362128 | 0.08016573 | 7.02896355 | 40.03122171 | 4/53 | PIK3CB, PIP4K2C, PLEKHA4, SYNJ2 |
| Formation of Intermediate Mesoderm | 0.003612934 | 0.080961121 | 28.51515152 | 160.3474045 | 2/8 | BMP4, FOXC2 |
| TGFBR3 Regulates TGF-beta Signaling | 0.003612934 | 0.080961121 | 28.51515152 | 160.3474045 | 2/8 | TGFB1, TGFBR2 |
| Attenuation Phase | 0.003713813 | 0.080961121 | 10.7298913 | 60.04121203 | 3/27 | HSPA6, COL4A6, TNFRSF21 |
| Activation of HOX Genes During Differentiation | 0.004243744 | 0.087512893 | 5.018614851 | 27.4132264 | 5/91 | JUN, POLR2E, HOXB3, RARB, HOXB2 |
| Activation of Anterior HOX Genes in Hindbrain Development During Early Embryogenesis | 0.004243744 | 0.087512893 | 5.018614851 | 27.4132264 | 5/91 | JUN, POLR2E, HOXB3, RARB, HOXB2 |
| Signaling by PDGF | 0.004656254 | 0.091277883 | 6.376516254 | 34.23898492 | 4/58 | COL4A5, PLAT, COL9A3, PIK3CB |
| Non-integrin membrane-ECM Interactions | 0.004950035 | 0.091277883 | 6.260262009 | 33.23172804 | 4/59 | LAMA5, TGFB1, COL4A6, COL4A5 |
| HSF1 Activation | 0.005024471 | 0.091277883 | 9.536231884 | 50.47942531 | 3/30 | HSPA6, COL4A6, TNFRSF21 |
| Laminin Interactions | 0.005024471 | 0.091277883 | 9.536231884 | 50.47942531 | 3/30 | LAMA5, COL4A6, COL4A5 |
| MET Activates PTK2 Signaling | 0.005024471 | 0.091277883 | 9.536231884 | 50.47942531 | 3/30 | LAMA5, HGF, MET |
| Nuclear Events (Kinase and Transcription Factor Activation) | 0.005574371 | 0.096951342 | 6.039990807 | 31.34499046 | 4/61 | EGR3, PPP2R1A, FOSB, TRIB1 |
| POU5F1 (OCT4), SOX2, NANOG Repress Genes Related to Differentiation | 0.005717969 | 0.096951342 | 21.38419913 | 110.4310322 | 2/10 | EOMES, HHEX |
| MET Receptor Recycling | 0.005717969 | 0.096951342 | 21.38419913 | 110.4310322 | 2/10 | HGF, MET |
| Developmental Biology | 0.005976373 | 0.09833932 | 1.776752649 | 9.096869468 | 27/1385 | SEMA5A, FOXC2, PIK3CB, HHEX, RELN, PERP, POLR2E, EOMES, IRX1, IRX2, EPHA7, JUN, MYOCD, SMAD3, TGFB1, JUP, CNTN6, TBX2, BMP4, PKP2, HOXB3, RARB, COL4A5, HOXB2, COL9A3, MET, STX1A |
| VEGFA-VEGFR2 Pathway | 0.006057599 | 0.09833932 | 4.589632325 | 23.43669018 | 5/99 | CYFIP2, JUP, THEM4, PIK3CB, SHB |
